# Systems Genetics of Single Nucleotide Polymorphisms at the Drosophila *Obp56h* Locus

**DOI:** 10.1101/2021.06.28.450219

**Authors:** Sneha S. Mokashi, Vijay Shankar, Joel A. Johnstun, Wen Huang, Trudy F. C. Mackay, Robert R. H. Anholt

**Affiliations:** Department of Genetics and Biochemistry and Center for Human Genetics, Clemson University, 114 Gregor Mendel Circle, Greenwood, SC 29646, USA; Department of Biological Sciences, Program in Genetics, North Carolina State University, Raleigh, NC 27695, USA; Department of Animal Science, Michigan State University, East Lansing, MI 48824, USA

**Author notes:** Joint corresponding authors, Address correspondence to Robert R. H. Anholt, or Trudy Mackay.

**Keywords:** quantitative genetics, genotype-phenotype map, pleiotropy, fitness traits, RNA sequencing, micro-environmental variation

## Abstract

Variation in quantitative traits arises from naturally segregating alleles with environmentally sensitive effects, but how individual variants in single genes affect the genotype-phenotype map and molecular phenotypes is not understood. We used CRISPR/*Cas9* germline gene editing to generate naturally occurring variants with different site classes and allele frequencies in the *Drosophila melanogaster Obp56h* gene in a common genetic background. Single base pair changes caused large allele-specific and sexually dimorphic effects on the mean and micro-environmental variance for multiple fitness-related traits and in the *Obp56h* co-regulated transcriptome. However, these alleles were not associated with quantitative traits in the Drosophila Genetic Reference Panel, suggesting that the small allelic effects observed in genome wide association studies may be an artifact of averaging variable context-dependent allelic effects over multiple genetic backgrounds. Thus, the traditional infinitesimal additive model does not reflect the underlying biology of quantitative traits.

## Main

Quantitative traits vary continuously in natural populations due to segregating alleles at many genes with environmentally sensitive effects^1,2^. Understanding the genetic and environmental basis of variation for quantitative traits is important for precision medicine, agriculture, and evolutionary biology. However, it is challenging to dissect the genotype-phenotype map at base pair resolution because quantitative trait locus mapping studies are limited in precision due to blocks of linkage disequilibrium (LD) in linkage and association mapping populations, within which molecular polymorphisms are not independent; and effects of individual rare variants cannot be evaluated using association mapping. In addition, most molecular polymorphisms associated with quantitative traits are in non-coding genomic regions and presumably affect complex organismal phenotypes via regulation of gene expression, not only of the gene most proximal to the variant, but also of co-regulated genes^3–6^. There is also a growing realization that naturally occurring polymorphisms can be associated with micro-environmental variance as well as mean values of quantitative traits; *i.e.*, the within-genotype phenotypic variance can differ between alternative alleles^7–15^.

Here, we used a CRISPR/*Cas9* mediated gene deletion and reinsertion strategy to generate an allelic series of closely linked single nucleotide polymorphisms (SNPs) in a 738 base pair region including the *Drosophila melanogaster Odorant binding protein 56h* (*Obp56h*) gene. *Obp56h* is an excellent candidate for CRISPR/*Cas9* germline gene editing since it is a member of a multigene family^16–18^ for which functional redundancy is likely to prevent lethality upon gene deletion; it is a small gene (651 bp) without nested genes; and the nearest genes are 12,891 bp upstream and 10,374 bp downstream. There is also evidence that *Obp* genes have pleiotropic effects on quantitative traits. Other members of the *Obp* gene family have pleiotropic functions that extend beyond their traditional roles in chemosensation^4,19–21^. RNA interference of *Obp56h* affects olfactory behavior^22^, avoidance of bitter tastants^23^, mating behavior^24^, and expression of co-regulated genes associated with lipid metabolism, immune/defense response, and heat stress^24^. *Obp56h* is expressed in chemosensory tissues^25^ and in the central brain^26,27^, which suggests functional pleiotropy at the *Obp56h* locus.

We generated an *Obp56h* null allele by inserting a transgene with a selectable marker in the endogenous *Obp56h* genomic location, and then excised the selectable marker and replaced it with transgenes containing the minor allele for each of five *Obp56h* SNPs that segregate in the *D. melanogaster* Genetic Reference Panel (DGRP)^28,29^, all in a common genetic background. Three of the SNPs are common, with minor allele frequencies (MAF) ≥ 0.05, and two are rare (MAF < 0.05); two are protein coding missense polymorphisms and three are potentially regulatory variants located upstream and downstream of the gene body and in the 3’ UTR (Fig. 1). We quantified the effects of each SNP on the mean and micro-environmental variance of multiple fitness-related quantitative traits and on the transcriptome. This enabled us to compare the pleiotropic effects of multiple SNPs in one gene that are in LD in a natural population and of rare *vs.* common, protein coding *vs.* noncoding variants on organismal quantitative traits and the co-regulated transcriptome. We found extensive functional pleiotropy of *Obp56h*, and heterogeneous, large, and sexually dimorphic allelic effects for all organismal and transcriptional phenotypes. This reverse genetic engineering strategy can be generally applied to other genes to dissect variation in the genotype-phenotype relationship at single base pair resolution.

**Figure 1.**
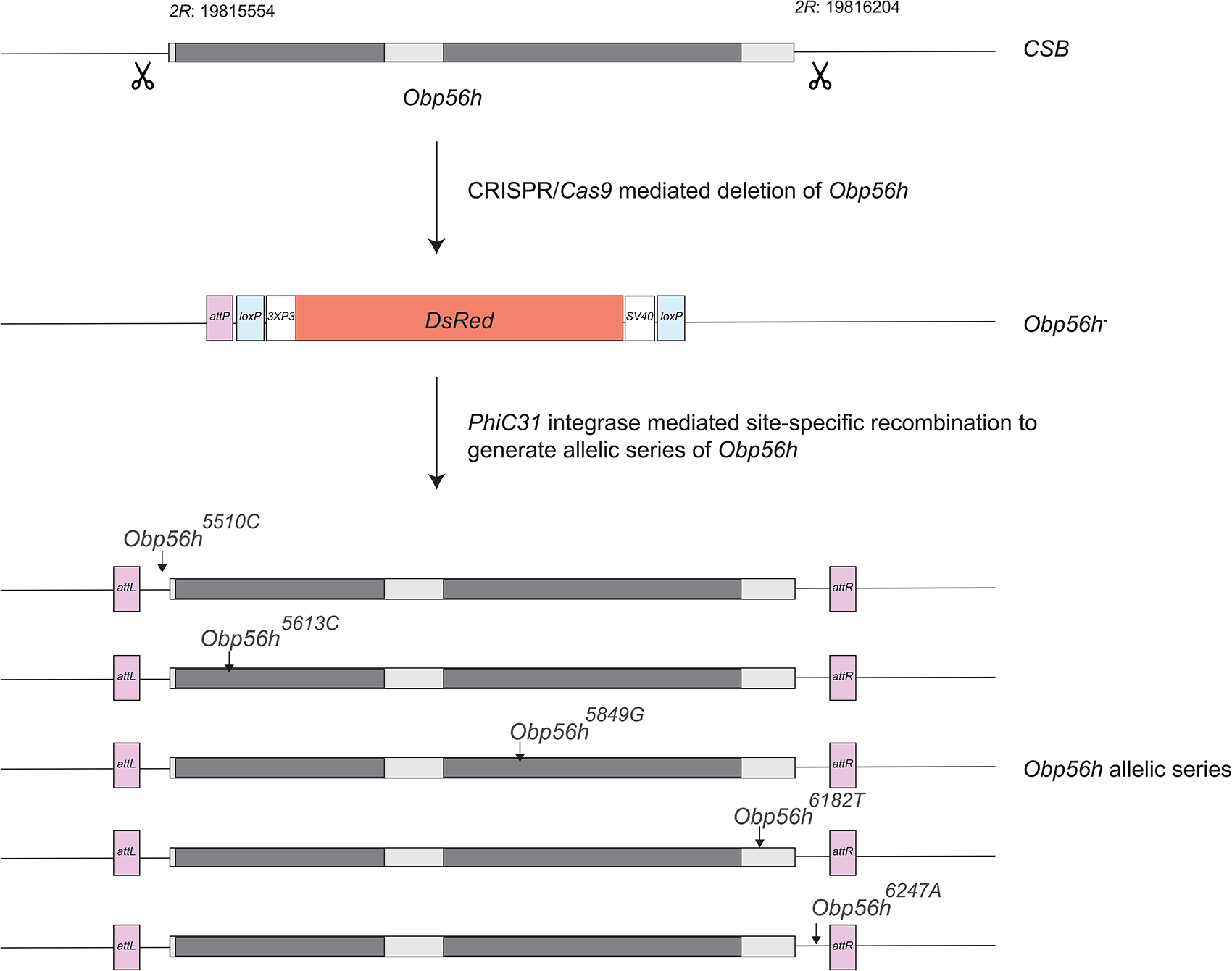
Construction of the *Obp56h* null allele and a series of DGRP minor alleles of *Obp56h*. Dark gray boxes represent exons of the *Obp56h* gene and light gray boxes indicate the intron and 5’ and 3’ untranslated sequences. We designed guide RNAs flanking the *Obp56h* gene for CRISPR/*Cas9-*mediated deletion at the cut sites, indicated by the scissor symbols, in the *Canton S-B* (*CSB*) genetic background. We replaced the gene with a cassette that contains a *DsRed* fluorescent marker (orange box) under the control of an eye-specific *3XP3* promoter and with SV40 polyadenylation sequences, *loxP* sites (blue boxes) for *Cre*-mediated removal of the insert, and an *attP* site (purple box) for *PhiC31*-mediated reinsertion. We then performed *PhiC31*-mediated site-specific recombination to generate *Obp56h* alleles with indicated nucleotide substitutions (arrows) that were generated by site directed mutagenesis. The *Obp56h* alleles are in the *CSB* background (which has the major allele for each of the five *Obp56h* SNPs) except for each single substituted base pair and the short 34 bp *attL* and 60 bp *attR* sequences (purple boxes) that remained after recombination.

## Results

### Generation of *Obp56h* allelic series

There are a total of 104 SNPs and 16 insertion/deletion polymorphisms in the 2,651 bp genomic region including the *Obp56h* locus and 1 kb up- and down-stream of this locus in the DGRP. We selected five SNPs with MAFs ranging from 0.006 – 0.26: *Obp56h^A5510C^* is 44bp upstream of the annotated transcription start site; the minor alleles of *Obp56h^T5613C^* and *Obp56h^C5849G^* aremissense mutations in the first and second exon, respectively; *Obp56h^A6182T^* is in the 3’ UTR; and *Obp56h^T6247A^* is 43 bp downstream of the annotated end site (Fig. 1; Table 1). The MAF of *Obp56h^C5849G^* and *Obp56h^A6182T^* are < 0.05 in the DGRP; these polymorphisms are underpowered for genome-wide association analyses. The SNP names begin with the common allele variant and end with the minor allele variant; the four intervening numbers are the last four digits of the genomic location. Allele names are the genomic locations followed by the nucleotide. Although LD declines rapidly with physical distance in *D. melanogaster*^28,29^, these SNPs are in strong LD in the DGRP and therefore not independent (Supplementary Table 1).

**Table 1.**
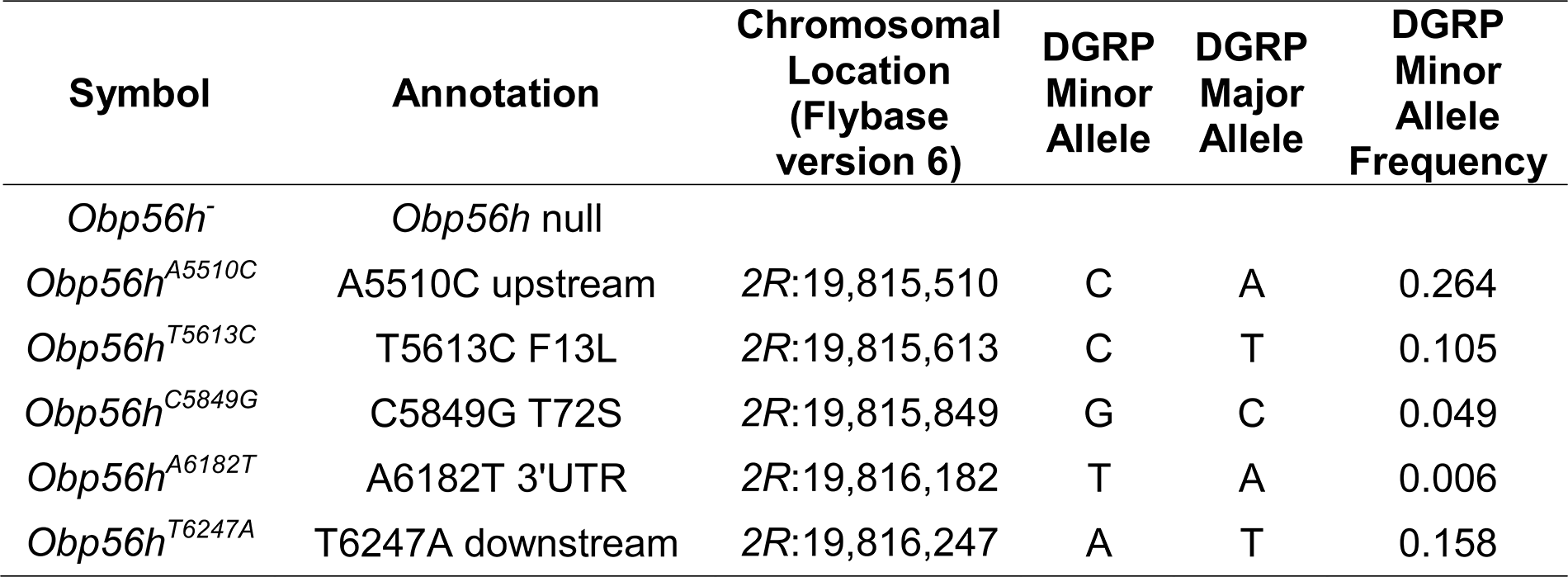
Genotypes used in this study. The control genotype, *CSB*, has the major allele for all *Obp56h* DGRP SNPs.

We designed a two-step strategy to assess the effects of individual SNPs on organismal phenotypes and the *Obp56h* co-regulated transcriptome. First, we used CRISPR/*Cas9* germline gene editing to generate a deletion of the *Obp56h* gene, substituting instead a *DsRed* fluorescent marker. This null allele is designated *Obp56h*^−^. In the second step, we inserted *Obp56h* genomic sequences that contain the minor allele of each of the five SNPs in the endogenous location to generate an *Obp56h* allelic series in a common genetic background (*CSB*, Fig. 1; Supplementary Fig. 1; Supplementary Table 2). The *CSB* allele has the common allele for each of the five SNPs. All transgenes were verified by Sanger sequencing.

### Effects of *Obp56h* alleles on organismal phenotypes

We assessed the effects of *Obp56h* null and SNP alleles on the mean values of several fitness-related traits: viability, sex ratio, feeding behavior, starvation stress resistance, time to recover from a chill-induced coma, heat stress resistance, locomotor activity, and sleep traits. All variants had reduced viability relative to the *CSB* control, with a greater effect for the SNP minor alleles (*P* < 0.0001) than the null allele (*P* < 0.05) (Fig. 2A; Supplementary Table 3). To assess whether effects on viability were different for male and female offspring, we calculated sex ratios and observed that the average number of eclosing males was less than the number of females for the *Obp56h^5613C^*, *Obp56h^5849G^*, *Obp56h^6182T^* and *Obp56h^6247A^* alleles (Fig. 2B; Supplementary Table 3).

**Figure 2.**
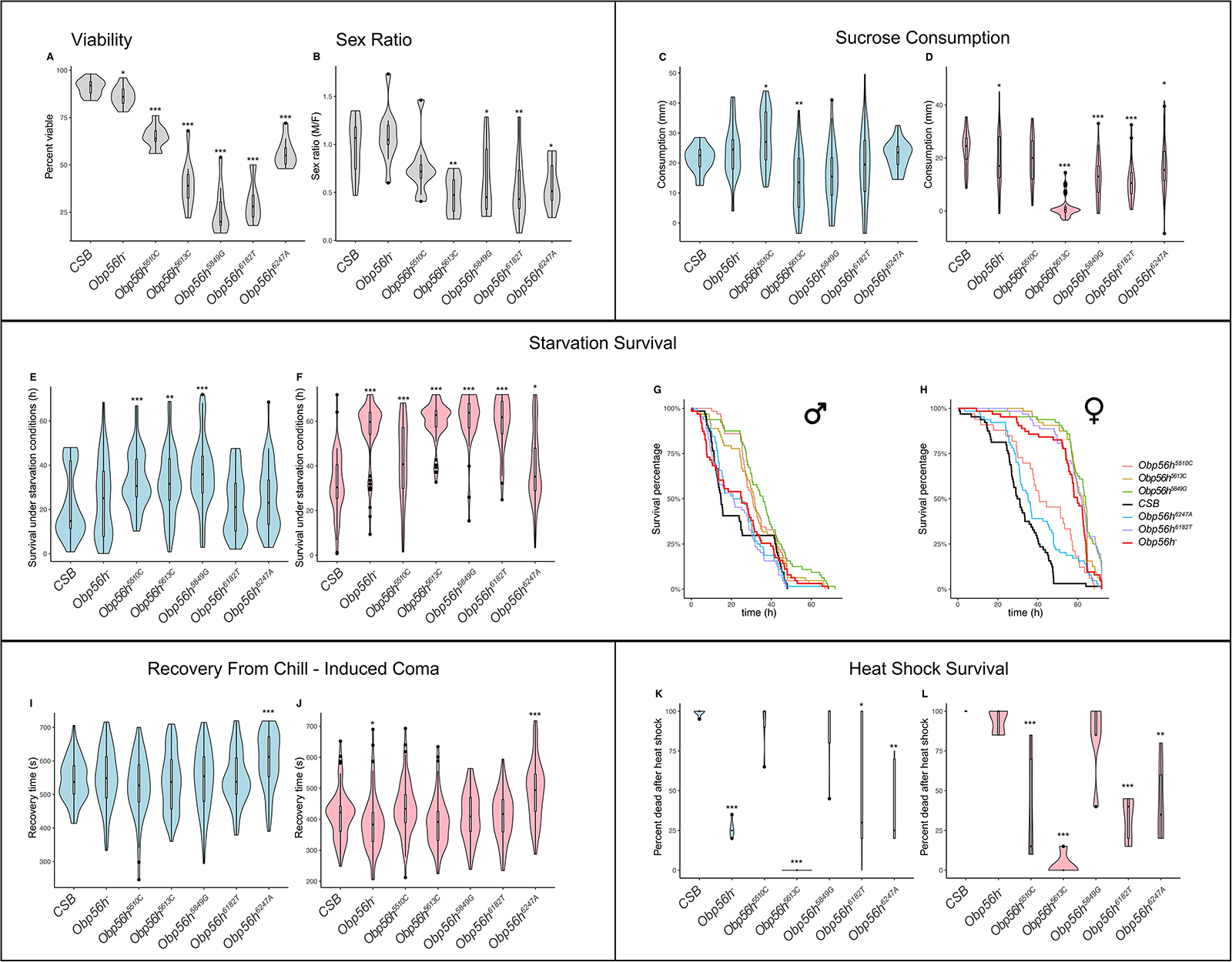
Pleiotropic effects of *Obp56h* alleles on fitness-related quantitative traits. (A) Viability. (B) Sex ratio. (C, D) Sucrose consumption. (E, F) Survival under starvation conditions. (G, H) Survival curves under starvation stress. (I, J) Recovery from a chill-induced coma. (K, L) Heat shock survival. For assays where males and females were scored separately, males are indicated in blue females in pink. *: *P* < 0.05; **: *P* < 0.001; ***: *P* < 0.0001 (Supplementary Table 3).

The *Obp56h* alleles had heterogeneous, often sexually dimorphic effects on all other quantitative traits. *Obp56h*^−^ females (but not males) consumed less sucrose than the *CSB* control (*P* < 0.05). The *Obp56h^5613C^* allele had the strongest effect on consumption levels, with both males (*P* = 0.0064) and females (*P* < 0.0001) drinking significantly less than the control (Fig. 2C and 2D; Supplementary Table 3). The *Obp56h^5510C^* allele had a male-specific increase in sucrose consumption (*P* = 0.029); and *Obp56h^5849G^* (*P* < 0.0001), *Obp56h^6182T^* (*P* < 0.0001) and *Obp56h^6247A^* (*P* = 0.012) had female-specific decreases in sucrose consumption (Fig. 2C and 2D; Supplementary Table 3).

*Obp56h*^−^ females (*P* < 0.0001), but not males, were more resistant to starvation stress than the control. All SNP minor alleles showed increased survival time under starvation conditions than the major allele in females. In males, alleles of *Obp56h^5510C^*, *Obp56h^5613C^* and *Obp56h^5849G^* had increased survival time under starvation stress; *Obp56h^6182T^* and *Obp56h^6247A^* were not significantly different from *CSB* (Fig. 2E, 2F, 2G, 2H; Supplementary Table 3). With respect to time to recovery from a chill-induced coma, the *Obp56h*^−^ allele slightly decreased recovery time (*i.e.*, in the direction of increased fitness) in females only (*P* = 0.03), while the only SNP to affect chill coma recovery time was *Obp56h^T6247A^*, for which the minor allele increased recovery time in females (*P* < 0.0001) and males (*P* = 0.0006) (Fig. 2I and 2J; Supplementary Table 3). The most heterogeneous effects of *Obp56h* alleles we observed were for survival following heat stress. *Obp56h*^−^ males (*P* < 0.0001), but not females, had increased survival compared to *CSB*. However, *Obp56h^5613C^* had markedly increased survival following heat stress in males (*P* < 0.0001) and females (*P* < 0.0001); *Obp56h^5849G^* was not significantly different from CSB in either sex; *Obp56h^5510C^* had a female-specific increase in survival time after heat stress (*P* = 0.0007); and *Obp56h^6182T^* and *Obp56h^6247A^* had increased survival times but with smaller effects than *Obp56h^5613C^* (Fig. 2K and 2L; Supplementary Table 3).

*Obp56h^5510C^* did not significantly affect total locomotor activity in either sex, but activity was increased in males for all other alleles and for *Obp56h^5613C^* and *Obp56h^5849G^* in females (Fig. 3A and 3B; Supplementary Table 3; the activity of *Obp56h*^−^ females decreased compared to *CSB* (Fig. 3B; Supplementary Table 3). In females, the proportion of time spent sleeping during the night was increased relative to *CSB* for *Obp56h*^−^ and minor alleles of all SNPs. Day sleep in females was similarly increased for all but *Obp56h^5510C^*, which was not significantly different from *CSB*. In contrast, only *Obp56h^6182T^* affected night sleep in males. *Obp56h*^−^ and *Obp56h^5510C^*, *Obp56h^5613C^* and *Obp56h^6247A^* had increased day sleep in males (Fig. 3C and 3D; Supplementary Table 3).

**Figure 3.**
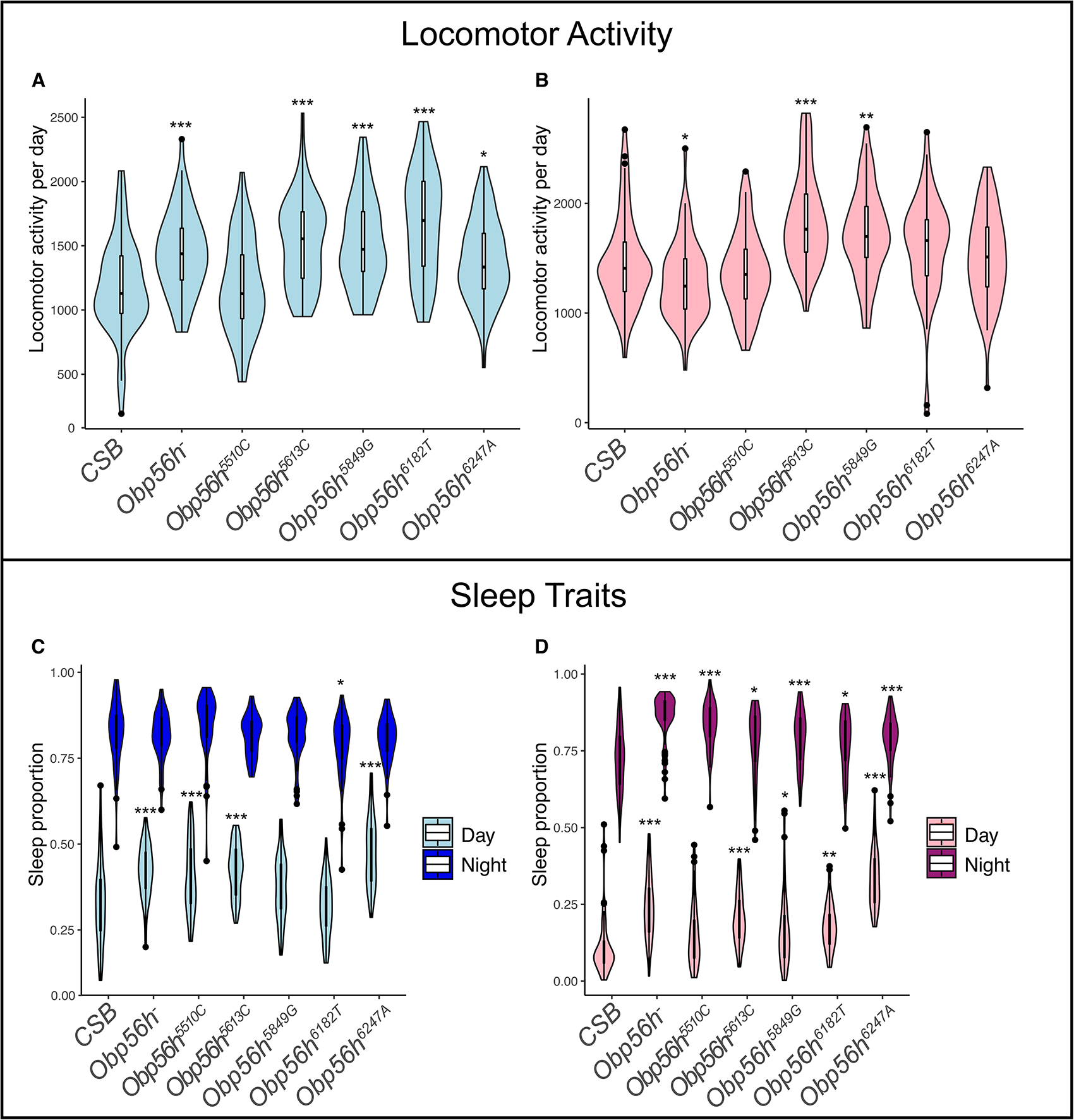
Effects of *Obp56h* alleles on activity and sleep phenotypes. (A, B) Total activity. (C, D) Sleep proportion during the day and night. Males are indicated in blue and females in pink. *: *P* < 0.05; **: *P* < 0.001; ***: *P* < 0.0001 (Supplementary Table 3).

In summary, single *Obp56h* SNPs have pleiotropic and sexually dimorphic effects on the mean values of all organismal quantitative traits we assessed (Supplementary Table 4). The effects of the common SNP and minor SNP alleles for each trait are heterogeneous, ranging from large to not significantly different from each other. Many of the alleles exhibit fitness trade-offs; for example, trade-offs between reduced viability and increased resistance to starvation and heat stress and between increased resistance to heat stress but a longer time to recover from chill coma stress. The SNP alleles typically have effects in the same direction as the null allele, but the SNP allele effects are often greater than the null allele effects.

### Effects of *Obp56h* SNPs in the DGRP

We previously evaluated the effects of the *Obp56h* SNPs for a subset of the organismal quantitative traits evaluated in this study using genome wide association analyses in the DGRP^11,12,29,30^. This affords us the opportunity to compare the effects of the same homozygous SNPs in a common genetic background *vs*. averaged over multiple genetic backgrounds. In contrast to the large and significant SNP effects observed in *CSB*, the effects were small and not significant in the DGRP (Supplementary Table 5). This observation is inconsistent with independent additive SNP effects and implies the existence of epistatic modifier loci in the DGRP that on average suppress the effects of the *Obp56h* SNPs on organismal phenotypes and/or variants in linkage disequilibrium with the *Obp56h* SNPs that counter their effects.

Effects of *Obp56h* alleles on micro-environmental variance of organismal phenotypes Micro-environmental variance (or general environmental variance^1^), refers to the phenotypic variation for a quantitative trait that occurs among individuals of the same genotype when they are reared in a common environment. We performed formal analyses of variance of micro-environmental variance for *Obp56h* alleles (Supplementary Table 4) and found that this phenomenon is pervasive: the micro-environmental variance for all alleles is significantly different from that of *CSB* for multiple organismal phenotypes. Changes in micro-environmental variance are allele-specific within each trait and are often sex-specific for each allele. The pleiotropic effects of *Obp56h* alleles on micro-environmental variance vary by trait and allele; *e.g.*, micro-environmental variance is largely increased for heat shock survival and largely decreased for sleep traits. Effects of *Obp56h* alleles on micro-environmental variance are decoupled from their effects on trait means: most alleles affect either the mean or the micro-environmental variance for any sex/trait combination, although some alleles affect both mean and the micro-environmental variance in the same or opposite directions for a given sex and trait (Supplementary Table 4).

### Effects of *Obp56h* alleles on genome wide gene expression

To investigate the cellular processes that might underlie the observed sexually dimorphic pleiotropic effects of *Obp56h* alleles, we obtained whole transcriptome profiles for heads from males and females separately, and identified differentially expressed genes among the *CSB*, *Obp56h*^−^ and *Obp56h* SNP minor alleles (Supplementary Tables 6 and 7). *Obp56h* expression is obliterated in both sexes in the *Obp56h*^−^ null allele compared to *CSB* and is partially restored in the reinsertion lines (Fig. 4A). At a false discovery rate (FDR) < 0.05, we identified 1,009 (717) differentially expressed genes in females (males) in any comparison between two alleles (Supplementary Table 7). A total of 406 co-regulated genes are in common between males and females, 603 are female-specific and 311 are male-specific. Gene set enrichment analyses^31^ reveal that differentially expressed genes in common between males and females and male-specific genes are enriched for terms involved in mitochondrial function, whereas genes that are only differentially expressed in females are enriched for terms involving protein translation, transport and localization, development and signal transduction (Supplementary Table 7).

**Figure 4.**
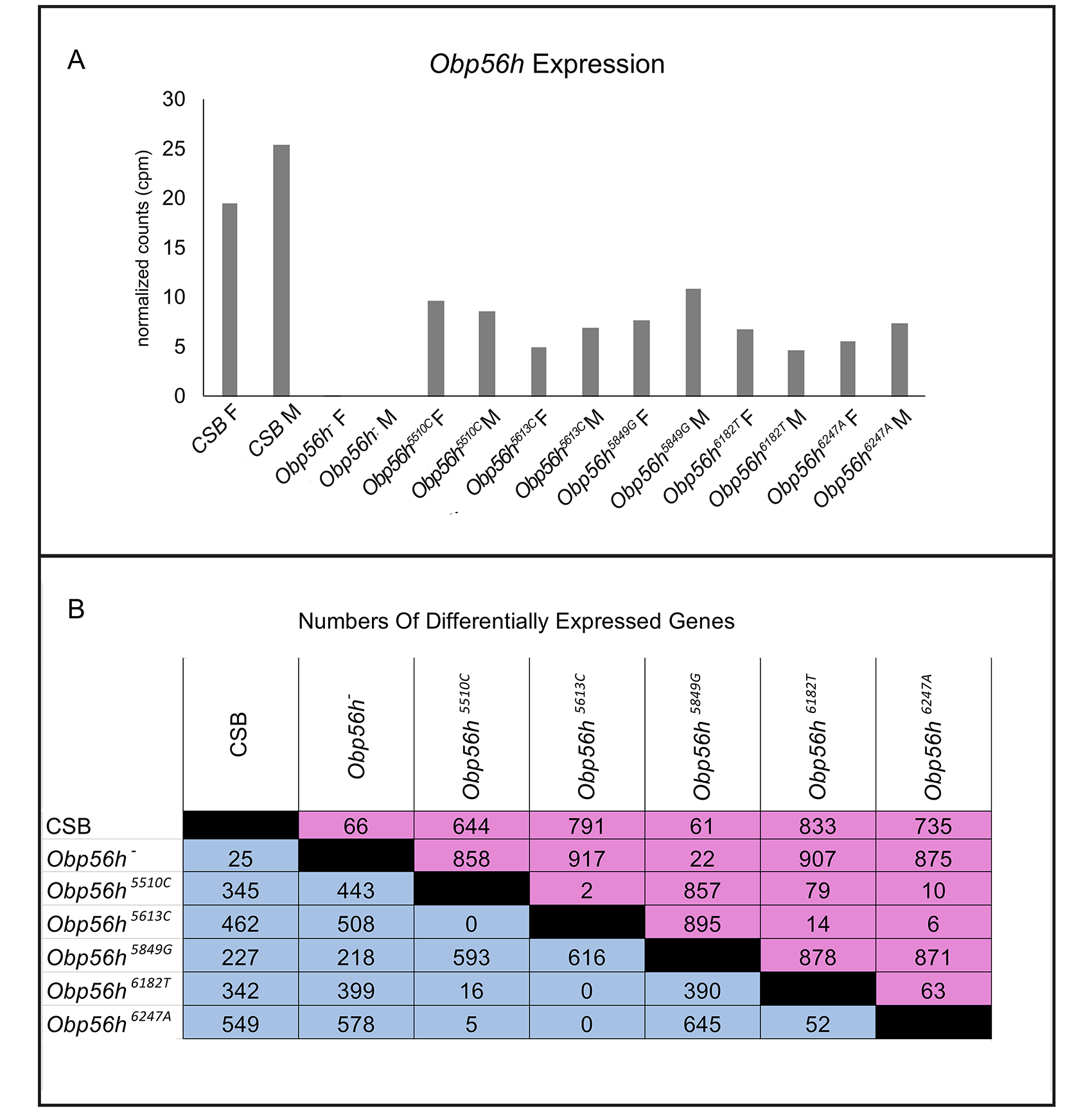
Summary of RNA sequencing analyses of *Obp56h* alleles. (A) Average normalized *Obp56h* expression counts for males (M) and females (F) for all genotypes. (B) Numbers of differentially expressed genes (FDR < 0.05) for every pairwise comparison of *Obp56h* alleles. Females (pink) are above the diagonal and males (blue) are below the diagonal.

Pairwise comparisons between the genotypes for females and males separately show that the number of coregulated differentially expressed genes varies greatly among alleles within each sex and between sexes for each pair of alleles (Fig 4B). However, in general the majority of the co-regulated genes have increased expression in *Obp56h^5510C^*, *Obp56h^5613C^*, *Obp56h^6182T^* and *Obp56h^6247A^* relative to *CSB* and *Obp56h*^−^ in both sexes; while the same genes have decreased expression in *Obp56h^5849G^* relative to *CSB* in males and females (Supplementary Table 8). This pattern is reversed for a second, smaller group of co-regulated genes (Supplementary Table 8).

We mapped the genes encoding differentially expressed transcripts onto known protein-protein interaction networks, separately for males (Fig. 5) and females (Fig. 6). The large network in each sex could be partitioned and clustered into smaller subnetworks that functionally converge toward oxidative phosphorylation, mitochondrial translation, circadian cycle, glutathione metabolism, ubiquitin-dependent proteolysis and cellular response to starvation in males (Fig. 5) and cytoplasmic translation, protein modification and localization, regulation of transport, G-protein coupled receptor signaling, mRNA splicing, chitin development and histone acetylation in females (Fig. 6). Both male and the female networks contained a large subnetwork that was enriched for electron transport chain and oxidative phosphorylation (Figs. 5 and 6).

**Figure 5.**
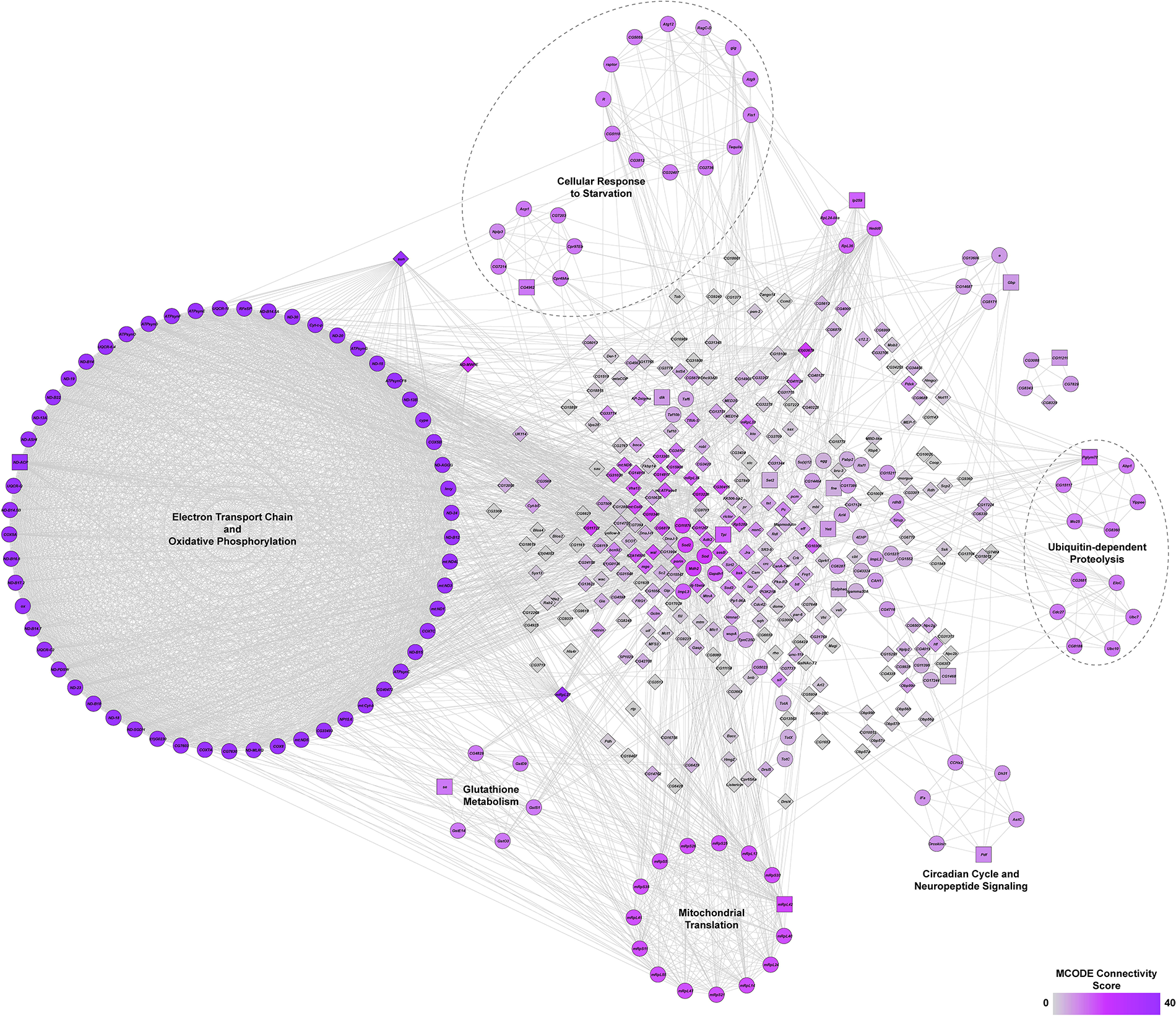
Protein-protein interaction network from differentially expressed transcripts among *Obp56h* alleles in males. The network was constructed using known interactions from the String database for all significantly (Benjamini-Hochberg FDR < 0.05) differentially expressed transcripts. Genes encoding the transcripts are organized into circular sub-networks based on MCODE clustering and the functional annotations of the sub-networks are based on statistically significant (Benjamini-Hochberg FDR < 0.05) enrichment of their Gene Ontology Pathways. The colors of the nodes represent the MCODE connectivity index and the shape of the nodes represents whether they are cluster seeds (squares), in cluster (circles) or unclustered (diamonds). Edges represent known protein-protein interactions.

**Figure 6.**
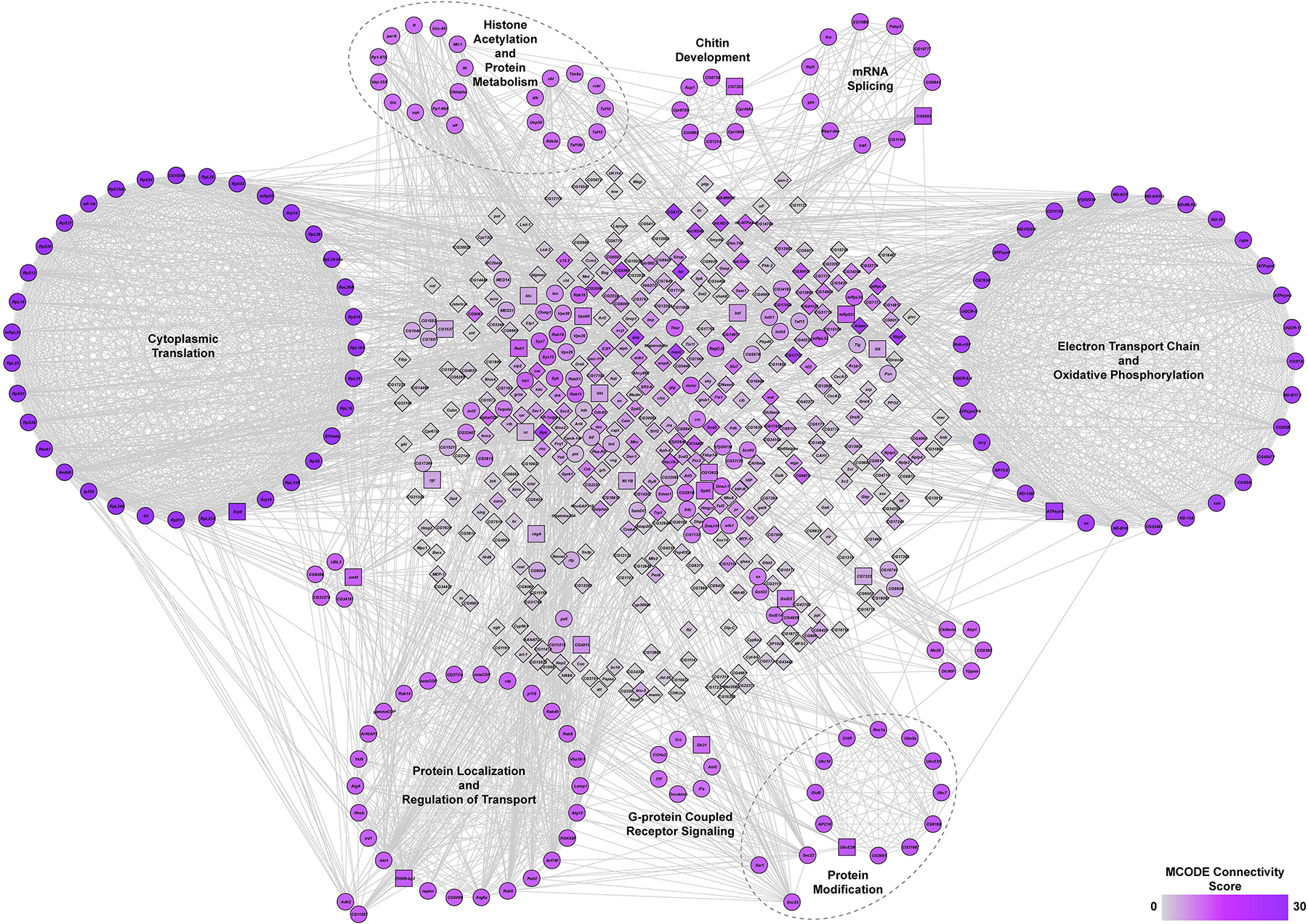
Protein-protein interaction network from differentially expressed transcripts among *Obp56h* alleles in females. The network was constructed using known interactions from the String database for all significantly (Benjamini-Hochberg FDR < 0.05) differentially expressed transcripts. Genes encoding the transcripts are organized into circular sub-networks based on MCODE clustering and the functional annotations of the sub-networks are based on statistically significant (Benjamini-Hochberg FDR < 0.05) enrichment of their Gene Ontology Pathways. The colors of the nodes represent the MCODE connectivity index and the shape of the nodes represents whether they are cluster seeds (squares), in cluster (circles) or unclustered (diamonds). Edges represent known protein-protein interactions.

### Effects of *Obp56h* alleles on micro-environmental variation of the transcriptome

We computed the coefficients of variation (*CV*, standard deviation/mean) across the replicates from normalized expression counts for each *Obp56h* SNP minor allele as well as for *CSB* and the *Obp56h*^−^ null allele. We plotted the distributions of *CV* across all expressed genes and all genotypes, separately in males and females (Supplementary Fig. 2). The distributions of *CV* were highly right-skewed; therefore, we chose genes with median expression above 10 counts per million across all alleles, for which at least one allele had a *CV* ≥ 1 and for which the variance heterogeneity analysis across alleles from Levene’s test had an FDR < 0.05 as contributing significantly to transcriptional micro-environmental plasticity. A total of 246 genes in males and 71 genes in females met these criteria for at least one variant. *Obp56h^6247A^* and *Obp56h^6182T^* had the largest number of transcripts with high micro-environmental plasticity in both sexes, and *Obp56h*^−^ had the smallest number of high plasticity transcripts in males and second smallest in females (Supplementary Table 9). Multi-dimensional scaling (MDS) based on the *CV* values showed that *Obp56h^6247A^* and *Obp56h^6182T^* were separated from the other alleles, but on different axes, indicating that the transcripts associated with high *CV* were different for these alleles (Supplementary Fig. 3). A total of 36 (50.7%) of the gene affecting micro-environmental plasticity of the transcriptome in females overlapped the genes associated with transcriptional micro-environmental plasticity in males. However, there was very little overlap between genes affecting the mean and micro-environmental plasticity of transcript abundances in either sex.

Gene Ontology (GO) enrichment analyses also showed distinct enrichment categories for *Obp56h^6247A^* and *Obp56h^6182T^*. Transcripts with high micro-environmental plasticity in *Obp56h^6247A^* males are enriched for terms involving immune response, response to stress, and lipase activity; whereas in *Obp56h^6182T^* males the enrichment was for transcripts associated with RNA binding, Box C/D RNP complex and the spliceosomal complex (Supplementary Table 9). Similar differences in enriched GO categories were observed in females but to a lesser extent due to the smaller number of transcripts with high micro-environmental plasticity (Supplementary Table 9). In addition to protein coding genes, regulatory non-coding transcripts contribute to micro-environmental variation in the transcriptome, especially snoRNAs and tRNAs (Supplementary Table 9).

## Discussion

The genetic architecture of quantitative traits inferred from many linkage mapping and genome wide association analyses in humans^32–34^ and model organisms, including *Drosophila*^35–37^, is highly polygenic, with large numbers of genes each with small additive effects, consistent with the infinitesimal model proposed by Fisher over a century ago^38^. The small effects could be because of imperfect LD between the genotyped variant and the true causal variant^32,39^, because the effects are truly small in all genetic backgrounds, or because the allelic effects are highly context-dependent and vary according to sex, environmental conditions and genetic background such that marginal (additive) effects over all contexts are small^40^. These possibilities can only be distinguished by examining the effects of naturally occurring SNPs in a common genetic background, which is now feasible using advanced germline gene editing technology^41,42^.

We found large effects of five naturally occurring SNPs in *Obp56h* on a battery of fitness-related organismal quantitative traits. The effects of the five SNPs varied in magnitude, and occasionally direction, within each trait, and the pattern of pleiotropic effects varied across traits. All alleles had sex-specific effects on at least one trait, but the pattern of sex-specificity was different for each trait/allele combination. There was no difference in the numbers of significant effects between rare and common variants (*P* = 0.64, Fisher’s Exact Test), as is typically found in genome wide association analyses^32,37^; nor between missense and regulatory variants (*P* = 0.49, Fisher’s Exact Test). We observed fitness trade-offs at the single variant level, which imposes an evolutionary constraint on natural selection at this locus (in the *CSB* genetic background). The observations that SNP allele effects are usually greater than those of the null allele and that the effects of the same alleles on the same traits measured in the DGRP were small and not significant are both consistent with genetic background effects (epistasis). The phenotypic effects of reduced expression of genes via RNA interference are often greater than those of null alleles^43–45^, thought to be due to a compensatory mechanism induced only by the null allele. Naturally occurring variants in the DGRP suppress the effects of new mutations^46,47^ and associations of DGRP alleles with quantitative traits vary according to population allele frequency, a hallmark of epistasis^24,40,48–50^. The naturally occurring *Obp56h* variants also affect the micro-environmental variance of multiple organismal quantitative traits. Together, all of these observations suggest that the small effects of alleles affecting quantitative traits in genome wide association studies are the consequence of averaging over multiple genetic backgrounds, males and females, and environmental contexts, and that effects in any one context may well be large. Although the infinitesimal statistical model fits these data, it obfuscates the underlying biology.

Most variants associated with quantitative trait variation are non-protein coding and could thus affect traits by perturbing expression of large genetic regulatory networks in relevant cell types, in the same way that a mutation in a single gene affecting a complex trait has quantitative effects on the abundance of many co-regulated transcripts^3,51,52^, called the transcriptional niche of the focal gene^6,21^. This concept is related to the omnigenic model of quantitative genetics^5^, which postulates that gene regulatory networks are highly interconnected, such that any variant in a core gene affecting a particular phenotype expressed in cell types relevant to the phenotype will affect many co-regulated genes. These concepts provide possible molecular bases for the highly polygenic, pleiotropic genetic architecture of quantitative traits.

This study provides support for these models for naturally occurring SNPs. Different variants in the *Obp56h* core gene have widespread *trans* effects on the transcriptome that are sex-specific and partially shared and partially distinct among the different alleles. Missense variants as well as variants in non-coding regions impact the *Obp56h* transcriptional niche; the largest number of co-regulated genes in both sexes is for *Obp56h^5849G^*, a rare missense variant. The enrichment of co-regulated genes involved in mitochondrial function provides a functional explanation for the sex-specific, pleiotropic effects of *Obp56h* variants on viability, food consumption, stress resistance, activity and sleep traits. Most *Obp56h* minor alleles affect increased transcription of mitochondrial genes, consistent with increased starvation and heat stress resistance, and increased activity and sleep duration. However, the correspondence between transcriptional coregulation and organismal phenotypes is not perfect. The *Obp56h^5849G^* allele has decreased expression of co-regulated genes relative to the other alleles, but the direction of the effects on organismal phenotypes is the same as for the other alleles, suggesting additional information than transcriptional co-regulation will be needed to predict effects on organismal phenotypes.

The *Obp56h* alleles also affect the micro-environmental plasticity of the transcriptome independent of the allelic effects on mean transcript abundance, providing a functional explanation for micro-environmental plasticity for organismal phenotypes. However, the *Obp56h^6182T^* and *Obp56h^6247A^* alleles, which have the largest number of transcripts with significant micro-environmental variance, are not different from the other alleles in terms of micro-environmental variance of organismal phenotypes, suggesting additional mechanisms buffer the transcriptome – organismal phenotype relationship.

We chose *Obp56h* for its favorable properties for CRISPR/*Cas9* gene editing and because previous studies suggested that this gene might have pleiotropic effects on the transcriptome and organismal phenotypes^22–24^. Members of the *Obp* gene family have been implicated in chemosensation as carriers of hydrophobic odorants^53,54^. In that context, there is differential expression of six other *Obp* genes (*Obp56g*, *Obp57a*, *Obp57b*, *Obp57c*, *Obp99b*, *Obp99c*) in males. Our results give further insight regarding the roles of *Obp* genes in additional non-chemosensory phenotypes^4,21,24^. *Obp56h* is expressed in the antenna and labellum^25^ and in cells of the central brain^26,27^. The ligand(s) for *Obp56h* in the brain are not known, but could be hydrophobic metabolites, which play a role in fundamental cellular processes that include mitochondrial metabolism and RNA processing. The extent to which naturally occurring polymorphisms affect these processes may lead to pleiotropic fitness phenotypes with different effects in males and females.

## Methods

### Generation of transgenic lines

The protocols used to generate the *Obp56h* deletion and allelic reinsertions are similar to those described previously^21,55,56^. Primer details are given in Supplementary Table 1. To generate a CRISPR/*Cas9* mediated null allele of *Obp56h* in a *CSB* genetic background we designed two guide RNAs flanking the gene using the Optimal Target Finder online tool^55^ and cloned them into the *pU6-Bbs1-chiRNA* plasmid. We used the *pBS-Hsp70-Cas9* plasmid as a source for *Cas9* and generated a donor plasmid containing *3XP3*-driven *DsRed* flanked by 1kb sequences homologous to the regions flanking the *Obp56h* gene. This vector also contains *loxP* sites flanking the *DsRed* cassette for subsequent removal of the cassette, and an *attB* site for site-specific *PhiC31* recombination to generate the reinsertion lines. We then generated the reinsertion alleles from the *Obp56h* deletion^21^. We generated allelic variants of *Obp56h* via site-directed mutagenesis in a *pattB* vector, which contained the *CSB* variant of the *Obp56h* gene. To generate an *Obp56h* allelic series, plasmids were injected into *Obp56h* knockout fly embryos (Model System Injections, Durham, NC).

### Fly husbandry

We reared all flies at 25°C, 60-75% relative humidity and 12-hr light-dark cycle on standard cornmeal-molasses-agar medium. Prior to experimentation, we reared the flies for two generations at controlled densities (5 males and 5 females per vial allowed to lay eggs for 2 days). We used 3-5-day old flies for all experiments.

### Organismal phenotypes

#### Viability and sex ratio

We placed 25 males and 25 females into egg collection cages with grape juice agar. We allowed the flies to acclimatize for 24 hr, with grape plate changes every 12 hr. After that, we changed the plates every 12 hr and collected 50 eggs per vial using a blunt moistened micro-probe and placed them in vials with standard culture medium. We scored the number and sex of flies that emerged until all pupae had eclosed from each of 10 vials per genotype.

#### Sucrose consumption

We performed capillary feeding (CAFÉ) assays as described previously^12,57^ with a single fly per vial. We scored a minimum of 18 flies per genotype and sex. *Starvation stress resistance:* We used *Drosophila* Activity Monitors to measure starvation stress resistance. We placed one fly per tube containing starvation medium (1.5% agar in distilled water) and ran the assay for 4 days in accordance with previous work^58^, with a total of 64 flies per sex per genotype. We obtained activity bout data using Shiny-R DAM^59^ and used the time of last activity bout as the time of death.

#### Recovery from chill-induced coma

We modified the original protocol for chill coma recovery assessment^60^ to enable us to measure accurately timepoints of recovery for 20 flies simultaneously by recording videos of the recovery period. For each genotype, we sorted 20 flies per vial, sexes separately, 4 replicates, into vials with 2ml food the evening before the assay. On the morning of the assay, we transferred the flies to empty vials and placed them in an ice bucket filled with wet ice for 3 hr. The ice-anesthesia 3-hr periods were staggered for the genotypes to be assessed on an assay day to allow us to record videos for approximately 30 minutes per vial. We gently placed the flies from the ice into wells of a 24-well microtiter plate with 2-5 flies per well for observation on an LED light box (Amazon) under a video camera (Canon). We recorded the flies for 30 min to determine how long it takes for each fly to right itself.

#### Response to heat shock

The day before measuring the response to heat shock, flies of each genotype were lightly anesthetized with CO_2_ and sorted in single-sex groups of 20 individuals in standard vials containing 5 ml food. On the day of the heat stress exposure, flies from each replicate vial were transferred without anesthesia into vials without food and placed in an incubator at 37°C (±0.5°C) for 180 min. After heat stress exposure, flies were immediately transferred to vials containing 5 ml of standard cornmeal-agar-molasses medium and returned to the 25°C incubator for 24 h. The percentage of surviving flies per vial was recorded 24h after the 3 hr heat shock. A fly was considered alive if it could move when the vial was gently tapped. We performed five replicates per genotype and sex.

#### Activity and sleep

We assessed total activity and proportion of sleep during the day and night^61,62^ using Drosophila Activity Monitors (TriKinetics). We ran the assay in accordance with previously published work^58^ and recorded data for 5 days on at least 64 flies per sex per genotype. We processed the initial data using Shiny-R DAM^59^.

#### Statistical analyses

For phenotypes for which measurements were obtained for both sexes, we assessed mean differences among the genotypes using factorial, fixed effects ANOVA models for all seven genotypes: *Y* = *μ* + *Genotype* + Sex + Genotype×*Sex* + *ɛ*, where *Y* is the phenotype, μ is the overall mean and *ɛ* is the residual (error) variance. For viability and sex ratio, we ran the reduced ANOVAs *Y* = *μ* + *Genotype* + *ɛ*. We also performed *t*-tests to identify the genotypes which were significantly different from the *CSB* control (planned comparisons). All analyses were performed using SAS Studio release 3.71 (SAS Institute, Cary, NC). To assess micro-environmental variance, we performed Levene’s and Brown-Forsythe tests of heterogeneity of within line variance, separately for males and females^13^ for all seven genotypes, and for pairwise comparisons between *Obp56h* alleles and CSB.

### RNA sequencing

To prepare libraries for RNA sequencing we collected 3-4 replicates of 50 flies, sexes separately, between 1pm and 3pm and flash froze them on dry ice in 15 ml Falcon tubes (Thermo Fisher Scientific, Waltham, MA). The flies were decapitated using a strainer (Carolina Biological Supply Company, Burlington, NC) for head collections^63^. The heads were collected on a dry ice-cooled fly pad and placed in 2ml pre-filled tough microfuge tubes with glass beads. Total RNA was extracted using the Direct-Zol microprep kit RNA extraction protocol (Zymo Research, Irvine, CA). The heads were homogenized in a bead mill (Thermofisher) for 1 min at 4m/s, after which the RNA was eluted with 15 μL water. We depleted ribosomal RNA using the NuQuant +UDI, Drosophila AnyDeplete kit (Tecan, Männedorf, Switzerland) and prepared bar-coded cDNA libraries for sequencing on an S1 flow cell on the NovaSeq 6000 platform (Illumina, San Diego, CA) as described previously^21^.

### Analysis of RNA sequences

We performed the initial steps of raw read processing and normalization of expression as previously described^21^. Briefly, we used the AfterQC pipeline^64^ to trim adapters, detect abnormal polynucleotide sequences, filter for low quality (Q□J<□J20) and short (<35 nt) sequence reads and generate of basic sequence quality metrics. We used the bbduk command from the BBTools package^65^ to detect rRNA contamination. We aligned high-quality sequence reads to the *Drosophila melanogaster* reference genome release 6 (version 6.13) using GSNAP aligner^66^ and mapped unique alignments to genes using the Subread package^67^. We excluded genes with fewer than 25% nonzero read counts or a median count of <2 from further analyses. We used GeTMM^68^ to normalize filtered expression counts. We ran ANOVAs across all seven genotypes (*Y* = *μ* + *Genotype* + *ɛ*) separately for males and females for each expressed transcript using PROC GLM in SAS Studio release 3.71 (SAS Institute, Cary, NC) to identify genes with significant (Benjamini–Hochberg FDR < 0.05) differential expression. We ran individual contrast statements for pairwise comparisons and then filtered them to only include genes that passed FDR in the overall model. We performed Gene Ontology analysis by statistical overrepresentation tests using PantherDB^31^. We generated protein-protein interaction networks from all differentially expressed genes (sexes separately) using the StringApp plugin of Cytoscape 3.8.2 followed by MCODE^69^ analysis to identify clusters of subnetworks. Functional annotation of the subnetworks was accomplished by performing Gene Ontology enrichment analysis on the membership. Labels were derived from GO biological processes with statistically significant enrichment (Benjamini–Hochberg FDR < 0.05).

Analysis of transcriptional micro-environmental plasticity was performed by first calculating the coefficient of variation (*CV*) across the replicates for each allele, separately for males and females, which showed that genes for which *CV* ≥ 1 were in the extreme right tail of the distribution. We also determined FDR values for Levene’s test of variance heterogeneity for estimates of between-replicate variance across all genotypes for each expressed gene, separately for males and females. Significant genes for transcriptional micro-environmental plasticity were those for which *CV* ≥ 1 for at least one allele, Levene’s test FDR for the gene < 0.05; and median normalized expression across all genotypes was 10 or greater counts per million. Multivariate ordination analysis was performed on the *CV* values of these genes for males and females separately using the *cmdscale* function that is part of the *stats* package in R. We also performed Gene Ontology enrichment analyses by allele and overall for co-regulated genes passing these criteria, separately for males and females.

## Acknowledgements

We thank Dr. Lakshmi Sunkara for help with RNA sequencing and Rachel C. Hannah for technical assistance with behavioral assays. This work was supported by NIH grant 1R01 GM128974 to TFCM and RRHA.

## Data availability

RNA sequence data have been deposited in the GEO data repository under accession number GSE178635. All code is available at https://github.com/snehamokashi/Systems_genetics_of_SNPs_at_Obp56h

## Author Contributions

SSM generated the *Obp56h* allelic series and performed all experiments; SSM and VS performed analyses on *Obp56h* differentially co-regulated transcripts; JAJ made *Obp56h*^−^; WH analyzed DGRP data; TFCM and RRHA conceptualized and directed the research program and provided resources; SSM, VS, TFCM and RRHA wrote the manuscript.

## Supplementary Material

**Supplementary Figure 1. Crossing scheme for the generation of the homozygous *Obp56h^−^* null allele and DGRP minor alleles for five DGRP SNPs (*Obp56h^mod^*) in the *CSB* genetic background.** All balancer and marker chromosomes and chromosomes for *PhiC31*-mediated insertion and *Cre*-mediated deletion were substituted into *CSB*. *w^1118^* denotes the *CSB X* chromosome. All third chromosomes are from *CSB* for all genotypes and are not shown.

**Supplementary Figure 2. Frequency distributions of *CV* between replicates for expressed genes for each genotype.** The most extreme *CV* values are for *CV* > 1 (vertical red line). (A) Males. (B) Females.

**Supplementary Figure 3. MDS plots of *Obp56h* alleles based on *CV* values from genes that contribute to micro-environmental plasticity of the transcriptome.** Multi-dimensional scaling plots represent the ordination of allelic variants, *CSB* and the *Obp56h*^−^ null allele based on the *CV* values for micro-environmental plasticity of the transcriptome. (A) Males. (B) Females. The percent overall variation explained by each axis is represented in the titles of the axes within parentheses.

**Supplementary Table 1. LD between *Obp56h* SNPs in the DGRP.** *Obp56h^A6182T^* is not included since the frequency of the minor allele is too rare to calculate LD in a sample of this size. *r*^2^ is above the diagonal and *D*’ is below the diagonal. All values of LD are significant at *P* < 0.0001 (χ^2^_1_ goodness of fit test)

**Supplementary Table 2. Primers used for generation and validation of *Obp56h* null and DGRP SNP minor alleles.**

**Supplementary Table 3. Effects of *Obp56h* alleles on organismal quantitative traits.** (A) ANOVA results for all genotypes. (B) Genotype means and significance of individual allele differences from *CSB* (*t*-tests).

**Supplementary Table 4. Summary of significance of differences of *Obp56h* alleles from *CSB*.** Entries in each cell are *P*-values. (A) Mean values. (B) Micro-environmental variance heterogeneity. MAF: Minor allele frequency. N/A: Not applicable.

**Supplementary Table 5. Comparison of effects and *P*-values for *Opb56h* alleles in the *CSB* genetic background and for the same alleles in the DGRP.** N/A: effect could not be estimated in the DGRP as the allele was not present in the sample of lines used to quantify chill coma recovery time. Data are from Ref. 11 (total activity) Ref. 29 (starvation survival, chill coma recovery); Ref. 12 (sucrose consumption); and Ref. 30 (viability).

**Supplementary Table 6. RNA sequencing raw data.** *Obp56h* alleles are denoted by their superscript. *CSB*: Canton S B control. M denotes males and F denotes females, and 1 and 2 indicate replicates 1 and 2, respectively. (A) Numbers of reads per gene. (B) Filtered and normalized counts/million reads. (C) Conditional means.

**Supplementary Table 7. ANOVA results of RNASeq data for seven *Obp56h* alleles.** Genes with significant (FDR < 0.05) variation among genotypes are shown. (A) Males. (B) Females. DF: Degrees of freedom. SS: Sums of Squares. MS: Mean Squares. F: F ratio statistic. (C) Genes in common between males and females and unique for each sex. (D) Gene set enrichment analyses.

**Supplementary Table 8. Significant (FDR < 0.05) differentially expressed genes in pairwise comparisons of *Obp56h* alleles.** log_2_FC is the log2 fold change of allele 2 relative to allele 1. Orange cells denote increased transcript abundance of allele 2 relative to allele 1, and purple cells represent decreased transcript abundance of allele 2 relative to allele 1. (A) Males. (B) Females.

**Supplementary Table 9. Micro-environmental plasticity for gene expression of *Obp56h* alleles.** (A) *CV* values for co-regulated genes in males. Entries above the threshold of *CV* = 1 are in bold font. Expression counts are the medians across all genotypes. *P*-Values and FDR are from Levene’s tests for variance heterogeneity across all genotypes. (B) Co-regulated genes with *CV* > 1 for each allele in males. (C) Gene Ontology (GO) enrichment for coregulated genes with *CV* > 1 in males. There was no enrichment for *Obp56h^5849G^* and *Obp56h*^−^. (D) *CV* values for co-regulated genes in females. Entries above the threshold of *CV* = 1 are in bold font. Expression counts are the medians across all genotypes. *P*-Values and FDR are from Levene’s tests for variance heterogeneity across all genotypes. (E) Co-regulated genes with *CV* > 1 for each allele in females. (F) Gene Ontology (GO) enrichment for co-regulated genes with *CV* > 1 in females. There was significant enrichment only for *Obp56h^5849G^* and *Obp56h^6182T^*.

