## Supplementary figures and images for "Systems Genetics of Single Nucleotide Polymorphisms at the Drosophila *Obp56h* Locus"

### Supplementary Figure 1

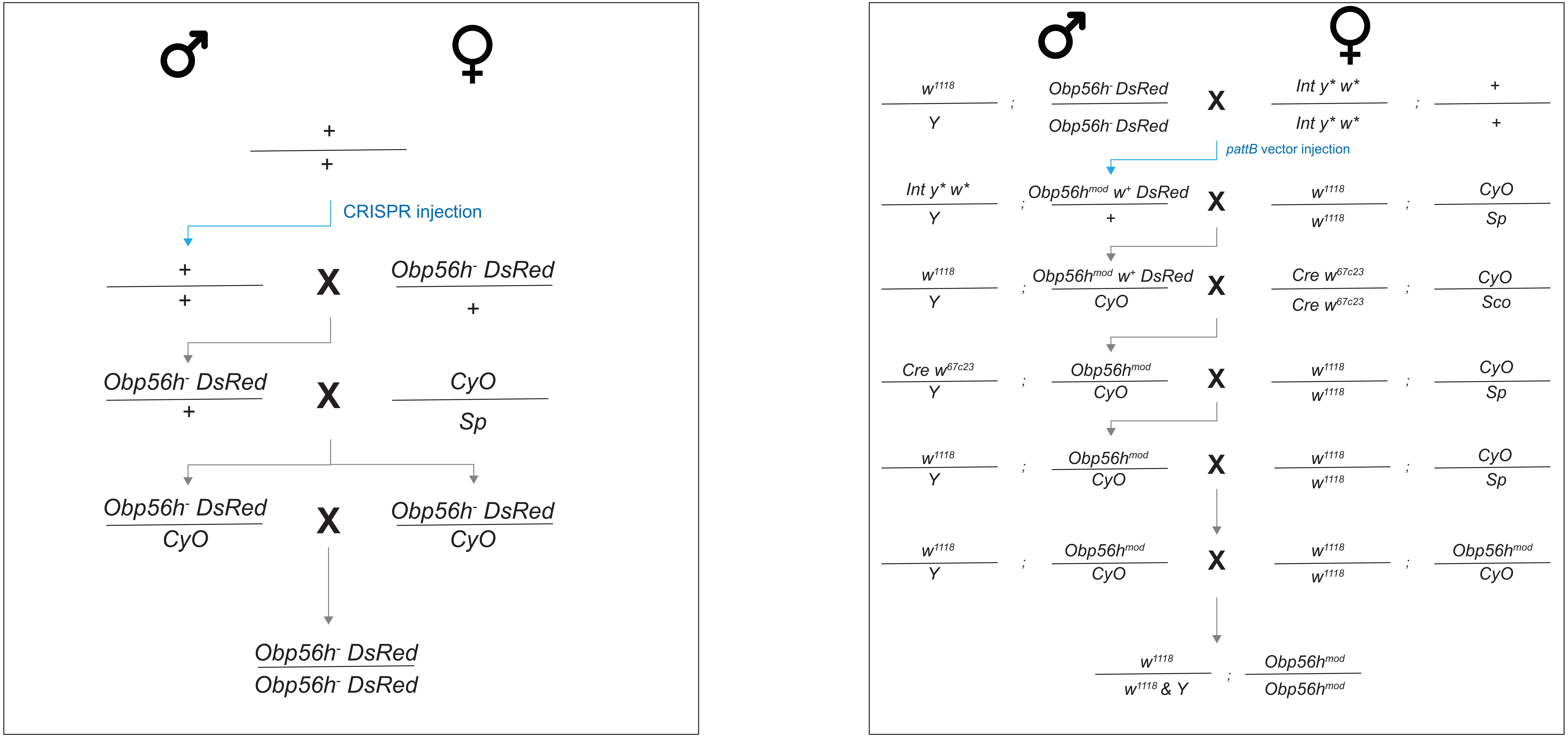

### Supplementary Figure 2

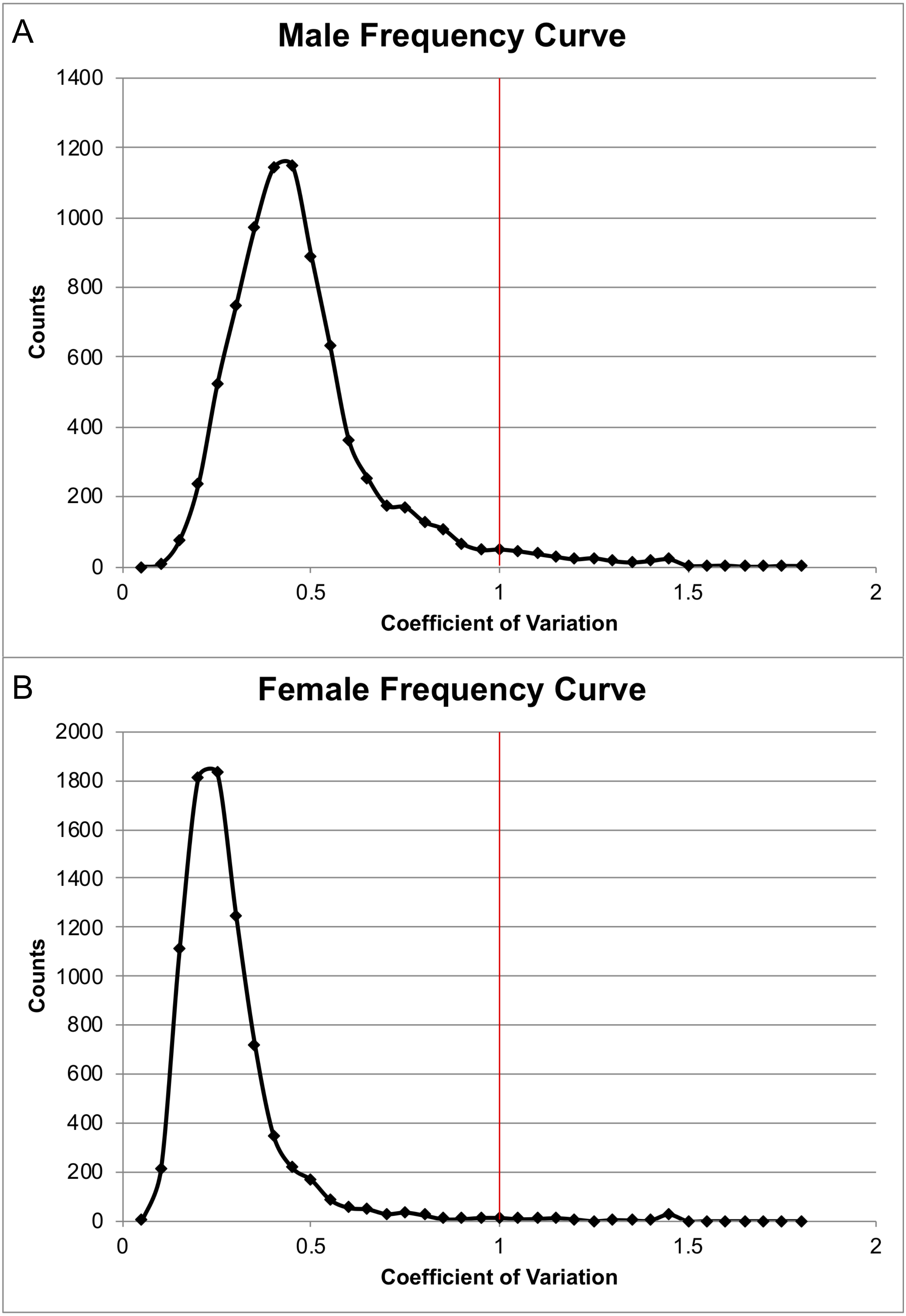

### Supplementary Figure 3

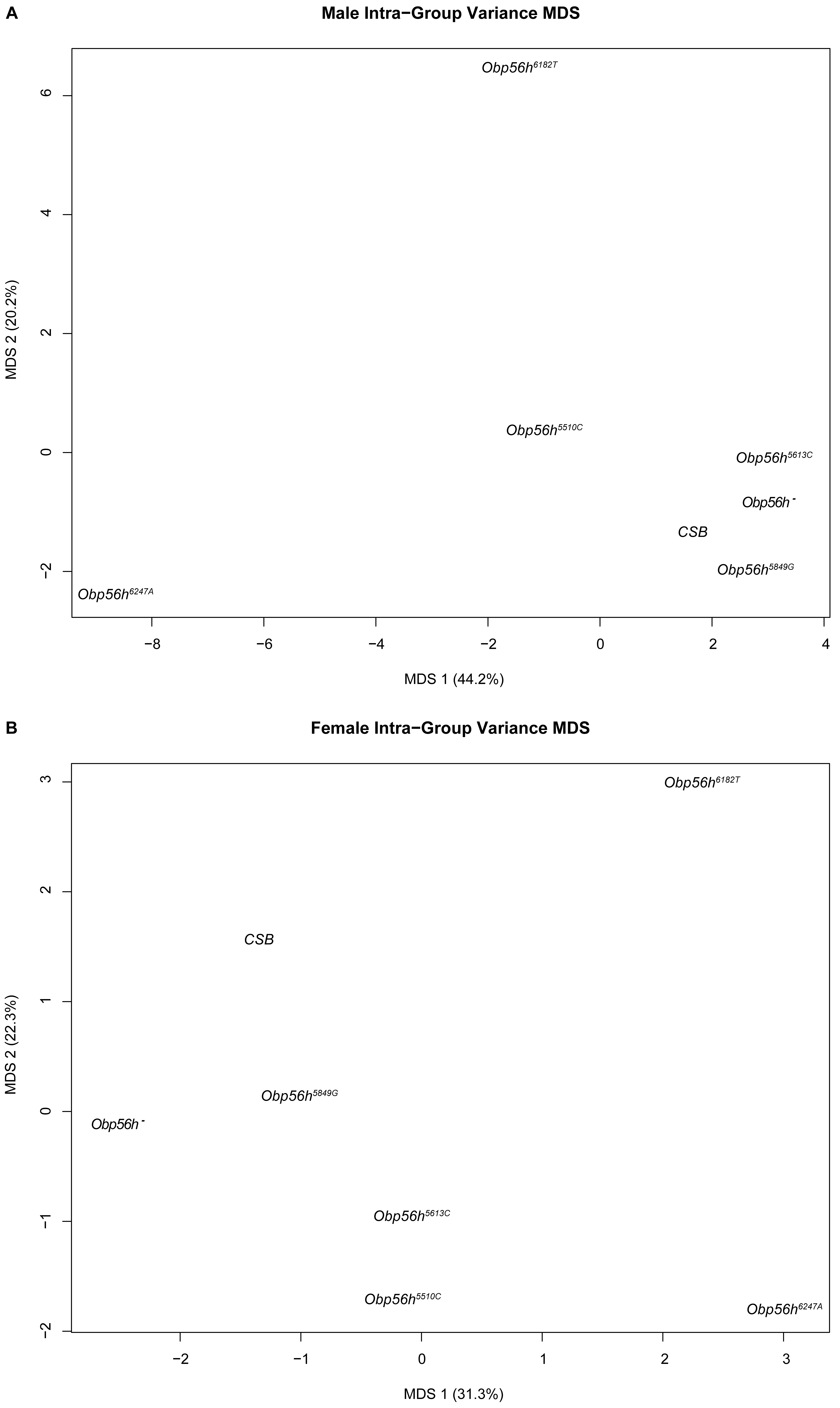
